# PatchWarp: Corrections of non-uniform image distortions in two-photon calcium imaging data by patchwork affine transformations

**DOI:** 10.1101/2021.11.10.468164

**Authors:** Ryoma Hattori, Takaki Komiyama

## Abstract

Two-photon microscopy has been widely used to record the activity of populations of individual neurons at high spatial resolution in behaving animals. The ability to perform imaging for an extended period of time allows the investigation of activity changes associated with behavioral states and learning. However, imaging often accompanies shifts of the imaging field, including rapid (∼100ms) translation and slow, spatially non-uniform distortion. To combat this issue and obtain a stable time series of the target structures, motion correction algorithms are commonly applied. However, typical motion correction algorithms are limited to full field translation of images and are unable to correct non-uniform distortions. Here, we developed a novel algorithm, PatchWarp, to robustly correct slow image distortion for calcium imaging data. PatchWarp is a two-step algorithm with rigid and non-rigid image registrations. To correct non-uniform image distortions, it splits the imaging field and estimates the best affine transformation matrix for each of the subfields. The distortion-corrected subfields are stitched together like a patchwork to reconstruct the distortion-corrected imaging field. We show that PatchWarp robustly corrects image distortions of calcium imaging data collected from various cortical areas through glass window or GRIN lens with a higher accuracy than existing non-rigid algorithms. Furthermore, it provides a fully automated method of registering images from different imaging sessions for longitudinal neural activity analyses. PatchWarp improves the quality of neural activity analyses and would be useful as a general approach to correct image distortions in a wide range of disciplines.

## Introduction

Two-photon microscopy has become a popular technique in neuroscience (Svoboda and Yasuda, 2006). It allows imaging of neuronal structures and activity inside the intact brain at high spatial resolution. The most common approach to measure neural activity with two-photon microscopy is calcium imaging. Calcium imaging is based on the principle that the intracellular concentration of calcium ion increases when neurons increase their firing rate. Therefore, calcium imaging allows an indirect measurement of neuronal spiking activity using fluorescent calcium indicators (Grienberger and Konnerth, 2012). Two-photon calcium imaging has several advantages over the classical techniques of recording neural activity using electrodes. For example, calcium imaging allows simultaneous recording of hundreds of neurons within a small area. It also allows longitudinal tracking of the activity of those neurons over weeks and months. Furthermore, specific cell types can be identified by targeting the expression of genetically encoded calcium indicators (GECIs) with viral or transgenic approach.

However, two-photon calcium imaging suffers from motion artifacts especially when it is applied to behaving animals. Such motion artifacts must be removed to accurately quantify neural activity. As such, a number of algorithms have been introduced to correct motion artifacts of two-photon calcium imaging data. Fast (∼100 ms) motion artifacts mostly reflect spatially uniform translations as long as each frame is scanned quickly within tens of miliseconds. These translations can be efficiently corrected by rigid alignment of individual image frames to a template image (Dubbs et al., 2016; Guizar-Sicairos et al., 2008; Mitani and Komiyama, 2018; Pachitariu et al., 2017; Pnevmatikakis and Giovannucci, 2017; Thévenaz et al., 1998). These rigid alignment methods have been extensively used for post hoc processing of two-photon calcium imaging data.

In recent years, it has become possible to perform calcium imaging for a long period of time, even for a continuous few hours without photobleaching due to improvement in calcium indicators (Chen et al., 2013; Dana et al., 2019). Improved stability and photosensitivity of calcium indicators allow stable calcium imaging with minimum laser power. Continuous long calcium imaging experiments are often necessary to understand the neural mechanisms of cognition, state representation, and learning (Hattori et al., 2017, 2019). However, rigid alignments of motion artifacts are often not sufficient for such long imaging sessions because the image fields can slowly distort over time. The slow image distortions can be caused by both biological and mechanical issues. For example, the shape of brain tissue may slowly change due to behaviors, metabolic activity, dilation of blood vessels, and liquid reward consumptions. Furthermore, mechanical instability of the animal stage or the microscope can also introduce slow non-uniform image distortions, particularly when the angle of the focus plane drifts over time. These image distortions limit the time of stable continuous imaging. Existing non-rigid motion correction algorithms (Giovannucci et al., 2019; Pachitariu et al., 2017; Pnevmatikakis and Giovannucci, 2017) partially mitigate this problem by splitting an imaging field-of-view (FOV) and estimating the optimal translational shifts for the multiple subfields, but such subfield-wise rigid translations are suboptimal to correct non-uniform distortions.

To overcome this limitation of two-photon calcium imaging, we developed a novel algorithm for the processing of two-photon calcium imaging data. The algorithm, PatchWarp, is a two-step algorithm with rigid corrections and warp corrections. First, PatchWarp corrects uniform motion artifacts by performing rigid alignments of image frames to a template image by iteratively re-estimating the template image. Then, it corrects image distortions by performing affine transformations on the subfields of each image frame. We show that PatchWarp effectively corrects image distortions of two-photon calcium imaging data of both cell bodies and axons, either through a glass window or an implanted gradient-Index (GRIN) lens. Furthermore, we show that PatchWarp can also register images from different imaging sessions for longitudinal across-session analyses of neural activity. We openly share the software at https://github.com/ryhattori/PatchWarp.

## Results

### Non-uniform slow image distortions during *in vivo* 2-photon calcium imaging

We first present example 2-photon calcium imaging sessions with non-uniform image distortions. We used a cell body imaging session (∼3.5 hours of imaging, retrosplenial cortex, camk2-tTA::tetO-GCaMP6s transgenic mouse (Mayford et al., 1996; Wekselblatt et al., 2016)) and an axon imaging session (∼20 minutes of imaging, primary motor cortex, AAV-hSyn-FLEX-axon-GCaMP6s (Broussard et al., 2018) in the basal forebrain of Chat-Cre transgenic mouse (Rossi et al., 2011)) as the examples. We compared the spatial positions of neuronal structures between early and late frames of each imaging session after correcting motion artifacts by rigid motion corrections (Figure 1). These examples clearly indicate the presence of non-uniform image distortions during imaging. Only some subfields are registered between early and late frames with a rigid motion correction algorithm. Corrections of these image distortions are of critical importance for the analysis of 2-photon calcium imaging data because a region-of-interest (ROI) drawn on a neuronal structure (e.g. cell body, axon bouton, spine) will be slowly displaced from the target structure in the presence of image distortions.

**Figure 1.**
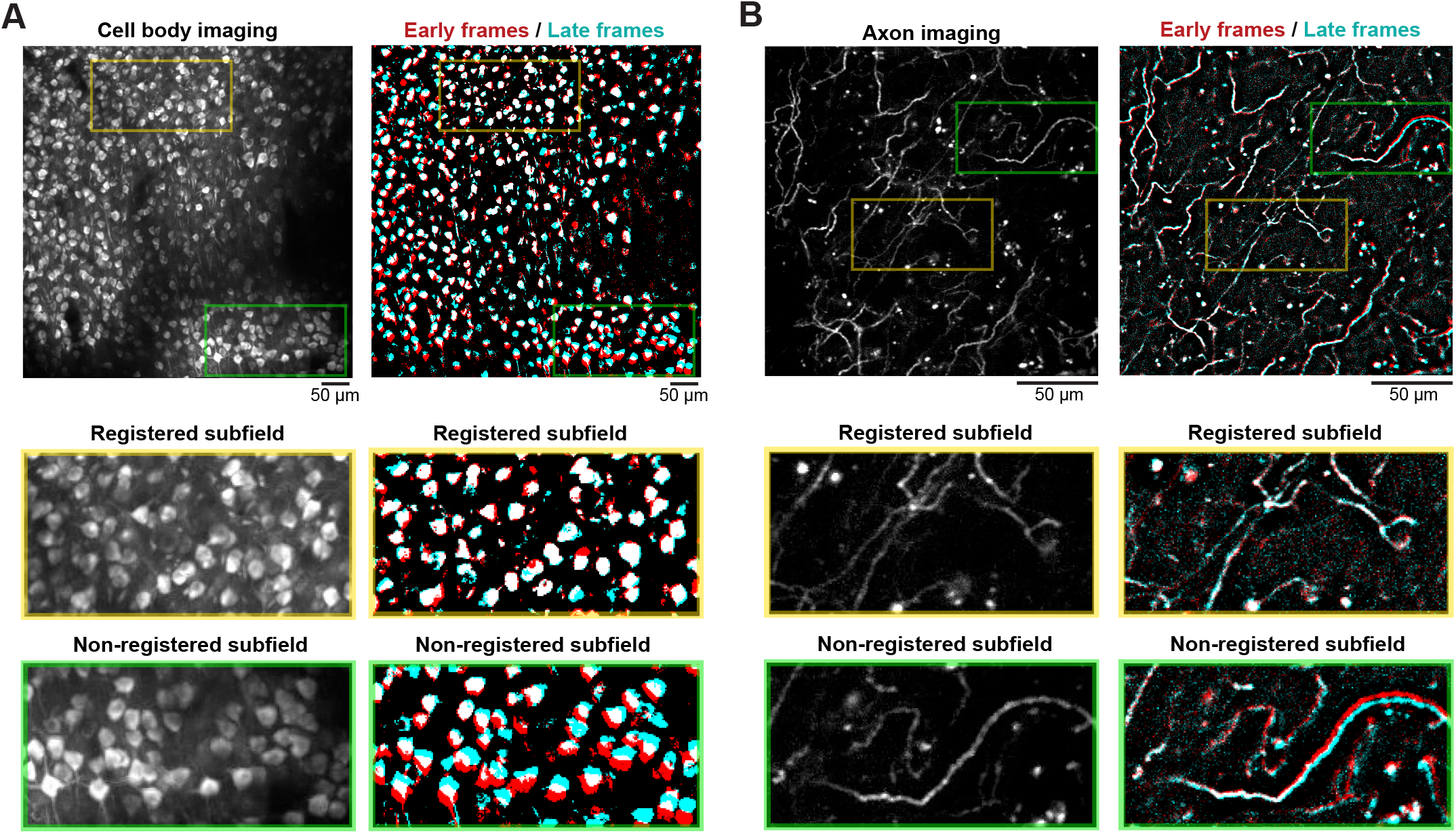
Non-uniform slow image distortions during *in vivo* 2-photon calcium imaging. **A)** Max-intensity projection image of an example RSC imaging session. The correct max-intensity projection after correcting both rigid moon artifacts and image distortions (*Left*), the max-intensity projections of the first (red) and the last (cyan) 14 min frames in the 3.5 hrs session without image distortion corrections (*Right*). The 2^nd^ and 3^rd^ rows show the zoomed images that highlight registered and non-registered subfields. The colors on the right panels are intentionally saturated with an arbitrary threshold to highlight the image overlaps. **B)** Max-intensity projection image of an example cholinergic axon imaging session. The correct max-intensity projection after correcting both rigid moon artifacts and image distortions (*Left*), the max-intensity projections of the first (red) and the last (cyan) 1.4 min frames in the 20 min session without image distortion corrections *(Right)*.

### PatchWarp algorithm for non-rigid image registration with distortion correction

We developed a novel algorithm, PatchWarp, to robustly correct both rigid translation and non-rigid distortion on 2-photon calcium imaging data. PatchWarp is a two-step algorithm which consists of rigid corrections and warp corrections (Figure 2A).

**Figure 2.**
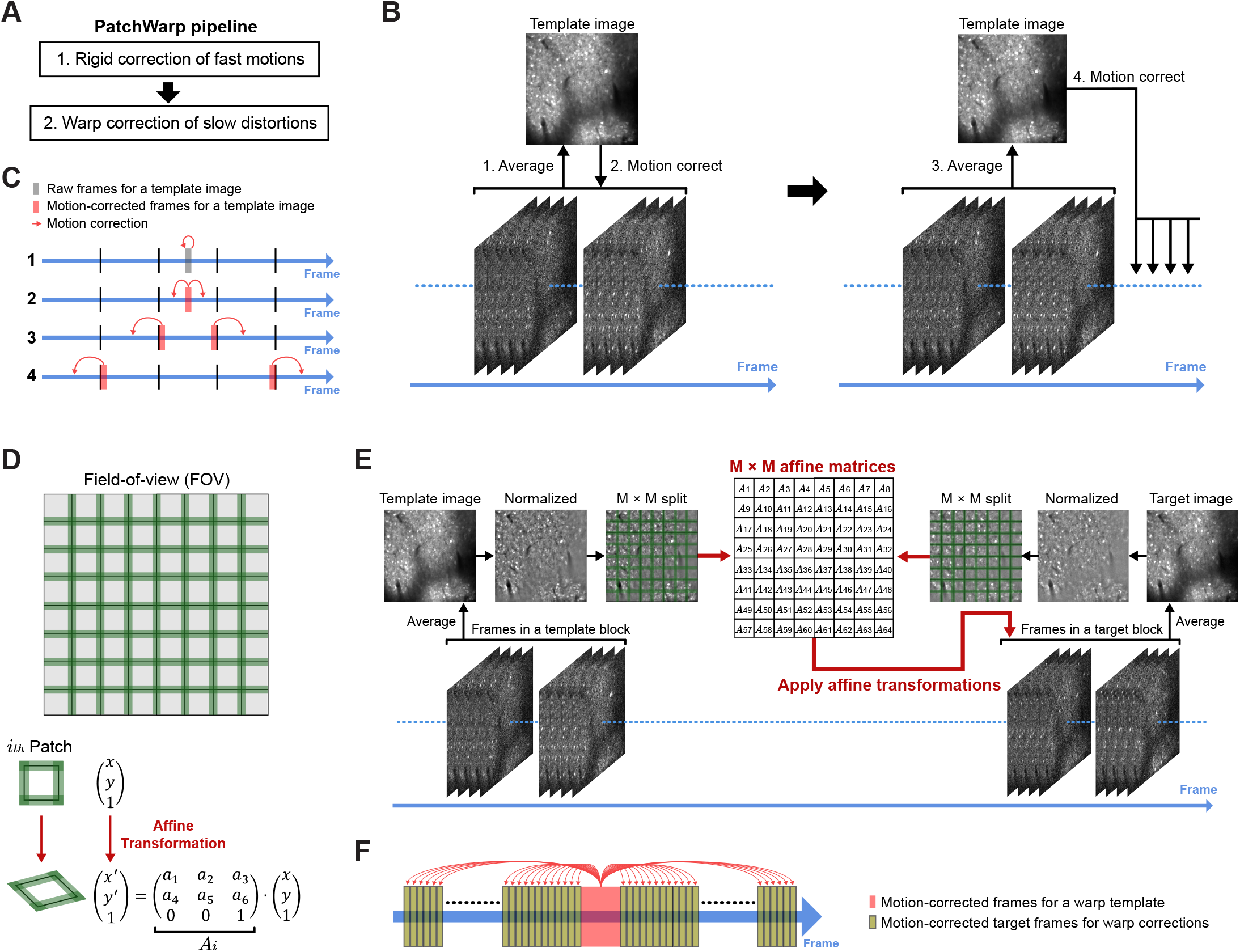
PatchWarp algorithm for correcting rigid moon artifacts and slow image distortions. **A)** PatchWarp is a two-step algorithm. The first step corrects the rigid moon artifacts, and the second step corrects the slow image distortions. **B)** Schematic of the rigid moon correction. The 1^st^ template image is created by averaging middle frames in a session, and these middle frames are registered to the mean image. The template image is updated by averaging the registered middle frames, and the updated template is used to register other frames. **C)** Timeline of the rigid moon correction. The rigid moon correction proceeds bi-directionally from the middle of a session. The template images are iteratively updated after specified frame number to reflect slow distortions on the template images. **D)** Warp correction is done by spliting a FOV into small overlapping patches and finding the optimal affine transformation matrix for each patch. Each patch is transformed by subpixel bilinear interpolation to fit the transformed coordinate. **E)** Schematic of the warp correction. Frames in the template and the target blocks are averaged, and the mean images are normalized by local intensity. The normalized images are split into patches, and an affine transformation that maximizes ECC between the template image and the transformed target image is estimated for each patch. The obtained transformations are applied to individual frames in the target block. **F)** Timeline of the warp correction. Warp correction uses a fixed template image that is created by averaging the middle frames of a session. Optimal affine transformations are independently estimated for each target block. The target block size should be determined based on the distortion speed in the imaging data.

The rigid motion correction was done by identifying the translational shift that maximizes the correlation between a template image and each image frame. We estimated the optimal translational shifts with subpixel accuracy using a pyramid-based hill climbing algorithm (Adelson et al., 1984; Mitani and Komiyama, 2018). We found that, when the net slow image distortions over the course of an imaging session are severe, a single template image is often not sufficient to accurately register all image frames because the difference between the single imaging template and the distorted frames becomes larger over time. Therefore, PatchWarp iteratively updates the template image during the rigid motion correction (Figure 2B and C). The 1^st^ template image is the average of the frames around the middle of a session. This template is used to motion correct the middle frames, and the template is updated by averaging the motion-corrected middle frames. Then, the motion correction proceeds bi-directionally from the middle towards both ends. Along the way, the template image is iteratively updated to reflect slow image distortions. This iterative template updating makes the rigid motion correction robust on imaging sessions with slow image distortions.

The rigid motion correction step is followed by a warp correction. As the examples in Figure 1 show, the image distortions on 2-photon calcium imaging data can be non-uniform across the field-of-view (FOV). This non-uniformity makes it impossible for a single geometric transformation function to fully register the images. Therefore, we implemented a ‘patchwork’ approach where a FOV is split into subfields and the optimal transformations are separately estimated for each subfield (Figure 2D). The number of subfields can be adjusted for each imaging condition based on its distortion pattern. We transformed the coordinate of each patch using affine transformation. Affine transformation is a geometric transformation that preserves collinearity and ratios of distances. It can transform an image by translation, rotation, scaling, and shearing. Affine transformation of an image is done by a matrix multiplication between the original image coordinate and an affine transformation matrix. The original image can be transformed to match the transformed coordinate by subpixel bilinear interpolation such that the image spatially fits the new coordinate. By using the optimal affine transformation matrix for each patch, we can correct non-uniform image distortions that exist in the original full-FOV. The optimal affine transformation matrix of each patch can be estimated by a gradient-based algorithm that aims at finding the optimal transformation matrix that maximizes the enhanced correlation coefficient (ECC) between a template image and an affine-transformed target image (Evangelidis and Psarakis, 2008; Psarakis and Evangelidis, 2005). We created the template image by averaging motion-corrected frames around the middle of a session (Figure 2E and F). In contrast to the rigid motion correction step, we use only a single template for all frames in a session for the second step of non-rigid warp correction. The source images to calculate the transformation matrix was created by downsampling all frames by non-overlapping moving averaging of 500 frames. Therefore, the total number of affine transformations that we need to estimate is [# of patches] × [# of downsampled frames].

One remaining issue is that different neurons within each patch change calcium signals differently across time (i.e. the relative brightness across different cells is dynamic), which can affect the correlation between the template and the target images. To make the estimations of optimal transformations robust against the variability of calcium levels across neurons, we normalized the intensity of each pixel by dividing the pixel value by the mean intensity of the nearby pixels (Figure 2E and Figure S1). This local intensity normalization allows the gradient-based algorithm to focus only on the cellular structures while ignoring calcium dynamics.

### Performance of PatchWarp algorithm

To examine the performance and general applicability of the PatchWarp algorithm, we used 3 different *in vivo* 2-photon calcium imaging conditions that were recorded from awake behaving mice. The first condition is the imaging of neuronal cell bodies from 6 different dorsal cortical areas (ALM: anterior-lateral motor area, pM2: posterior secondary motor cortex, RSC: retrosplenial cortex, PPC: posterior parietal cortex, S1: primary somatosensory cortex, V1: primary visual cortex, n = 1 session from each area, camk2-tTA::tetO-GCaMP6s transgenic mice) through glass windows. The second condition is the imaging of neuronal cell bodies from a ventral cortical area (OFC: orbitofrontal cortex, n = 6, camk2-tTA::tetO-GCaMP6s transgenic mice) through gradient-Index (GRIN) lenses which were implanted in the brains. The third condition is the imaging of axons from a dorsal cortex (M1: axons of basal forebrain cholinergic neurons imaged in the primary motor cortex, n = 6, AAV-hSyn-FLEX-axon-GCaMP6s in Chat-Cre transgenic mice) through glass windows.

We used 2 metrics to quantify the registration performance, ‘*Mean max-intensity difference (mMD)*’ and ‘*Mean correlation with mean image (mCM)*’. To calculate mMD, the maximum projection image across all frames is calculated for pre- and post-registered images separately. mMD is the difference in the means of all intensity values across pixels between these two images (mMD = post-mean – pre-mean). The mean of max-projection intensities across pixels takes a larger value in the presence of motion artifacts because movement of a neuronal structure across frames increases the total number of pixels to which the neuronal structure with calcium signals contributes in the max-projection image. Therefore, negative mMD indicates reduced motion artifacts after registration. Note that we downsampled frames by 50-frame non-overlapping moving averaging before creating the max-projection images to suppress the contribution of image noise to the max-projection intensity. The second metric, mCM, is the mean correlation between individual frames and a mean image. mCM increases after successful registration because motion artifacts reduce the similarity between individual frames and the mean image. The mean image to which individual frames are compared for the calculation of correlation coefficient can be either the mean of pre- and post-registered frames. Therefore, we also defined self-mCM as the CM when the used mean image was created by averaging the source frames of the mean image, and cross-mCM as the mCM when the used mean image was created by averaging the frames of the other compared condition (e.g. correlation between individual pre-registered frames and the mean image of post-registered frames).

First, we examined the performance of the rigid motion correction step. In all imaging sessions across the 3 different imaging conditions, mMD was consistently negative (Figure 3A). Furthermore, both self-mCM (Figure 3B) and cross-mCM (Figure S2) also consistently improved after registration. Therefore, rigid motion correction step significantly reduced motion artifacts in all imaging sessions (Supplementary video 1).

**Figure 3.**
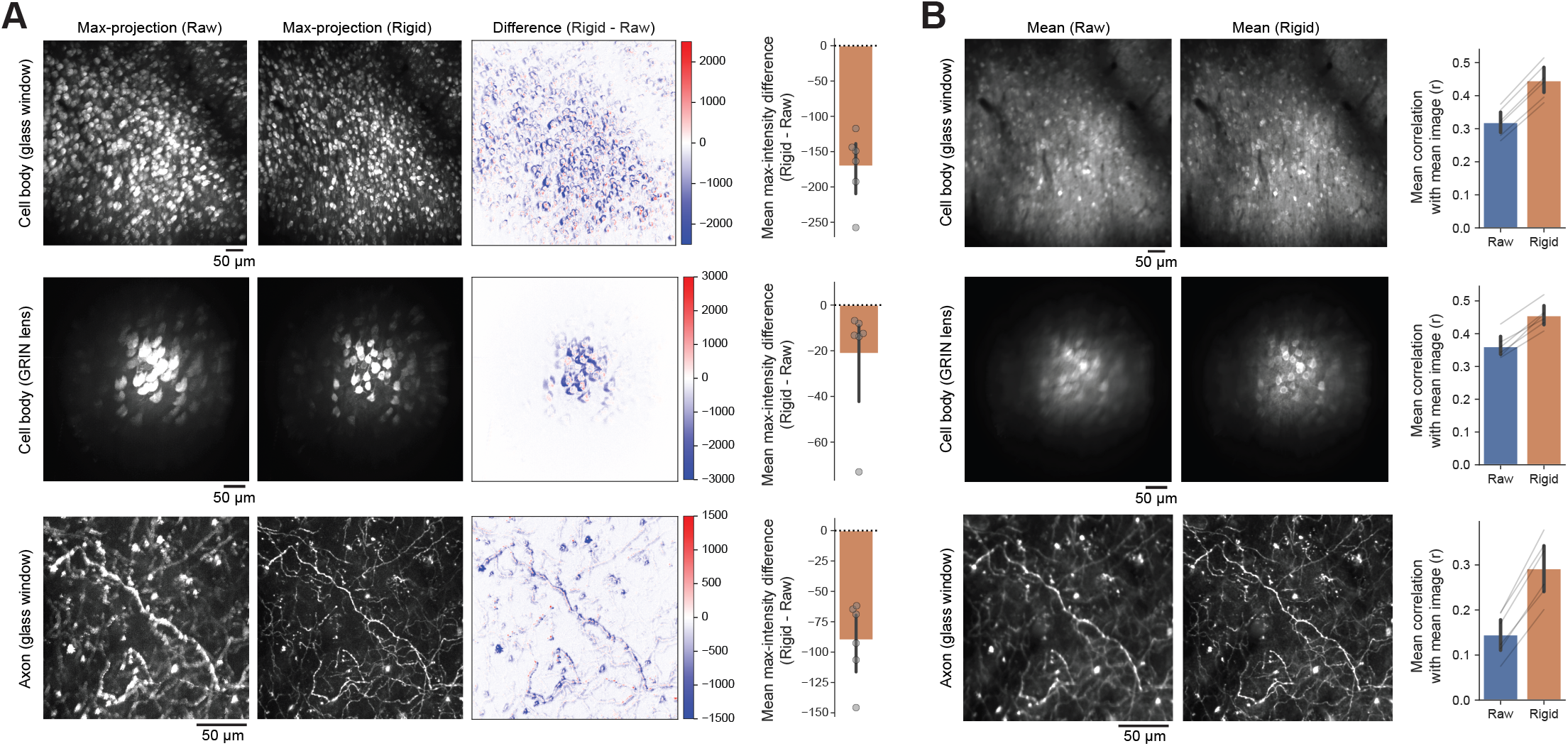
Performance of rigid corrections of moon artifacts. **A)** Max-intensity difference between pre- and post-rigid moon corrections. Max-projection images of raw frames, moon-corrected frames, and their intensity difference are shown. The bar graph indicates the mean max-intensity difference (mMD) within each FOV. mMD was negative in all sessions, indicating that moon artifacts consistently decreased after rigid moon corrections. **B)** Mean images of raw frames, moon-corrected frames, and the mean correlations between the mean images and individual frames of each condition (self-mCM). Self-mCM increased in all sessions, indicating that moon artifacts consistently decreased after rigid moon corrections.

Next, we examined the performance of the warp correction step. We compared mMD and mCM metrics between images after the rigid motion correction step and images after both the rigid motion and warp correction steps. We found that mMD (two-step mean – rigid mean) was consistently negative in all sessions (Figure 4A). Furthermore, both self-mCM (Figure 4B) and cross-mCM (Figure S3) consistently improved in all sessions. These results indicate that slow image distortions are ubiquitous in *in vivo* 2-photon calcium imaging data from awake behaving mice, and PatchWarp successfully corrects the distortions (Supplementary video 2). The consistent improvement across 8 different cortical areas (ALM, pM2, RSC, PPC, S1, V1, OFC, M1) with 3 different imaging conditions (cell bodies or axons through glass windows, cell bodies through GRIN lenses) indicate the general applicability of PatchWarp algorithm on 2-photon calcium data.

**Figure 4.**
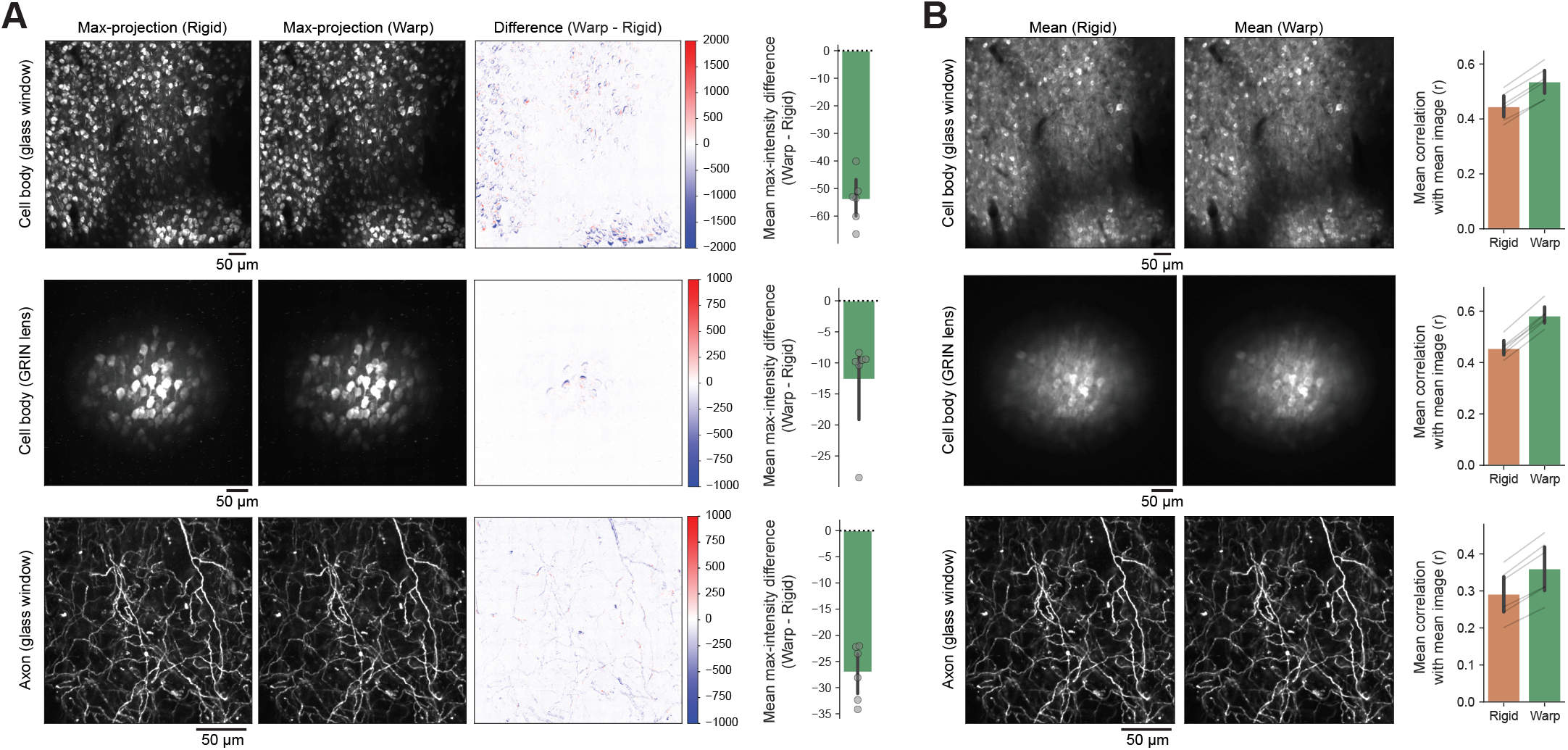
Performance of warp corrections of slow image distortions. **A)** Max-intensity difference between images with only rigid moon corrections and images with both rigid and warp corrections. Max-projection images of frames with only rigid moon corrections, frames with both rigid and warp corrections, and their intensity difference are shown. The bar graph indicates the mean max-intensity difference (mMD) within each FOV. mMD was negative in all sessions, indicating that warp corrections consistently improved the frame-by-frame registrations. **B)** Mean images of frames with only rigid moon corrections, frames with both rigid and warp corrections, and the mean correlations between the mean images and individual frames of each condition (self-mCM). Self-mCM increased in all sessions, indicating that warp corrections consistently improved the frame-by-frame registrations.

### Performance comparison to existing non-rigid registration algorithms

We compared the performance of PatchWarp to other existing non-rigid image registration algorithms. Suite2p (Pachitariu et al., 2017) and CaImAn (Giovannucci et al., 2019) are two of the most popular software for the processing of calcium imaging data. Both toolboxes offer non-rigid motion correction functions. The non-rigid motion correction algorithm in CaImAn is called NoRMCorre (Pnevmatikakis and Giovannucci, 2017), but we call it CaImAn here for simplicity. Non-rigid registration algorithms in Suite2p and CaImAn split a FOV into patches as in the warp correction step of PatchWarp. However, both Suite2p and CaImAn estimate only the translational shift of each patch. In contrast, the warp correction step of PatchWarp estimates affine transformations which can transform each patch by not only translation but also rotation, scaling, and shearing. Therefore, theoretically, PathWarp can correct more complex distortions than Suite2p and CaImAn. Furthermore, Suite2p and CaImAn use cross-correlation or phase-correlation to estimate the best translation while PatchWarp uses ECC. ECC has the property that it is invariant to changes in bias, gain, brightness and contrast (Evangelidis and Psarakis, 2008). The use of ECC, along with the local intensity normalization (Figure S1), may also improve the registration performance.

We compared mMD and mCM metrics of images after their non-rigid registrations. We used Python versions of Suite2p (ver. 0.9.0) and CaImAn (ver. 1.9.3) for this comparison. To make a consistent comparison, we used the same number of patches and the same patch size for each imaging session across the 3 algorithms. We found that mMD relative to PatchWarp was posive on most images from Suite2p and CaImAn (Figure 5A). Furthermore, self-mCM was consistently highest with PatchWarp in all imaging sessions. Thus, at least in the datasets tested here, PatchWarp outperforms the commonly used algorithms for non-rigid moon correction.

**Figure 5.**
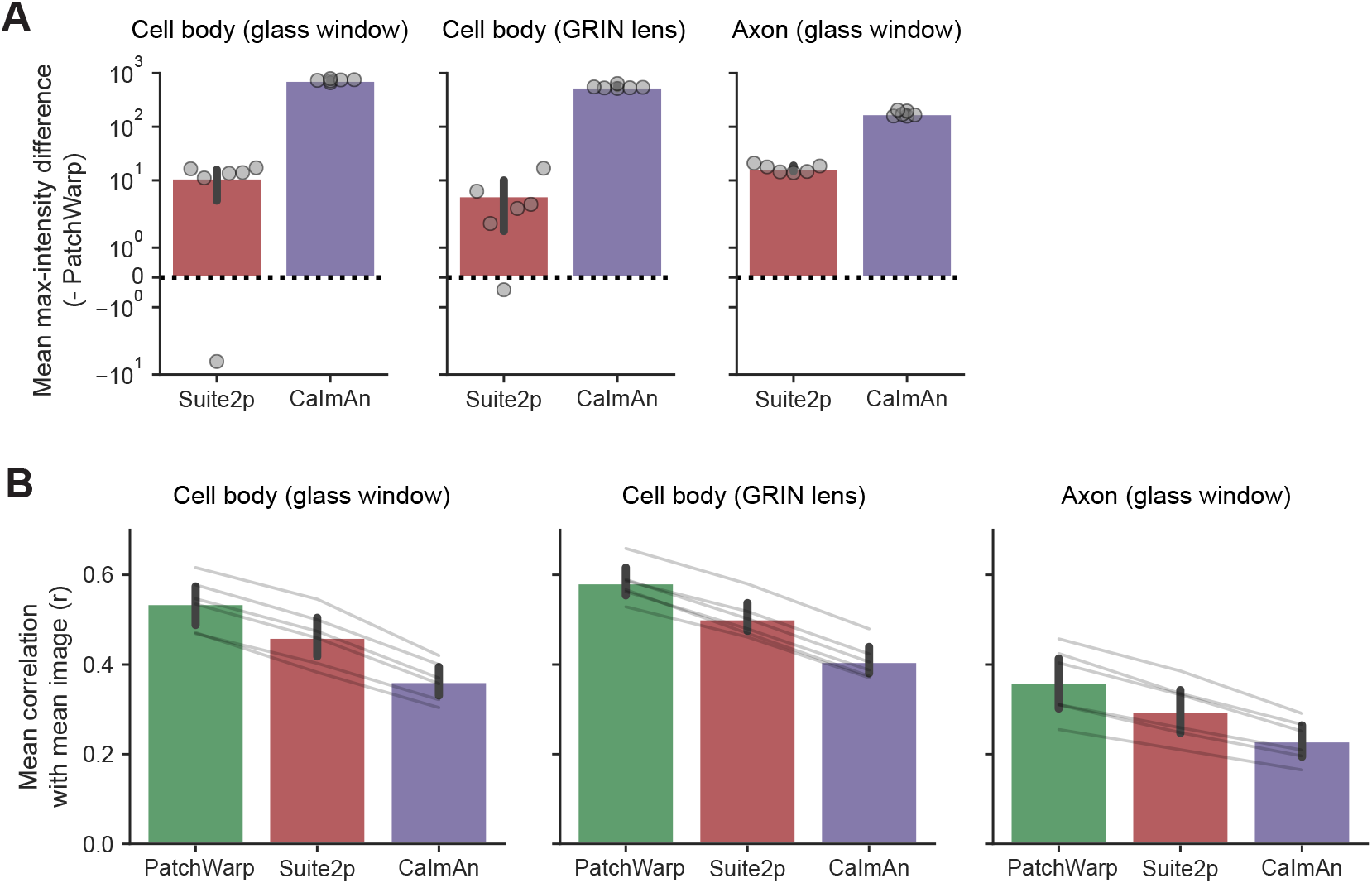
Performance comparison to existing non-rigid registration algorithms. **A)** Mean max-intensity difference (mMD) between images with PatchWarp registrations and images with non-rigid registrations of either Suite2p or CaImAn. mMD was posive in all sessions for both Suite2p and CaImAn, indicating that registration accuracy was consistently better with PatchWarp. **B)** Mean correlations between the mean images and individual frames (self-mCM) for images that were processed by PatchWarp or non-rigid registration algorithms of Suite2p and CaImAn. Self-mCM was highest with PatchWarp, indicating that registration accuracy was consistently beer with PatchWarp.

### PatchWarp algorithm for fully automatic across-session registration

One of the major advantages of 2-photon calcium imaging over electrode recordings of neuronal activity is that we can reliably track the identical population of neurons across many days. On each day, experimenters try to find the consistent focal plane by adjusting the position of an objective lens. However, some variability in the position of the focal plane across days is inevitable (e.g. x-y-z coordinate, rotation angle of the FOV, rotation angle of the objective lens). In particular, the difference in the rotation angle of the objective lens introduces non-uniform image distortion between images from different imaging sessions. Furthermore, the relative positions of neuronal structures in a FOV can also slowly change across days. These non-uniform image distortions necessitate an image distortion correction algorithm to optimally register different imaging sessions for longitudinal neural activity analyses. We devised a two-step PatchWarp method to estimate the optimal transformation that can be used to transform images from a session to the coordinate of another session. Unlike the within-session registration that uses individual frames, the inputs to the algorithm for across-session registration are summary images (e.g. mean image, max-projection image) from each imaging session. As the demonstration, we used 2 RSC imaging sessions that were acquired on different days and estimated the transformation that optimally transforms the session H image to the session G coordinate (Figure 6). We artificially made the registration problem harder than the original condition by adding shifts (translation and rotation) and barrel distortion to demonstrate the robustness of our algorithm even in the presence of large displacements and complex distortion.

**Figure 6.**
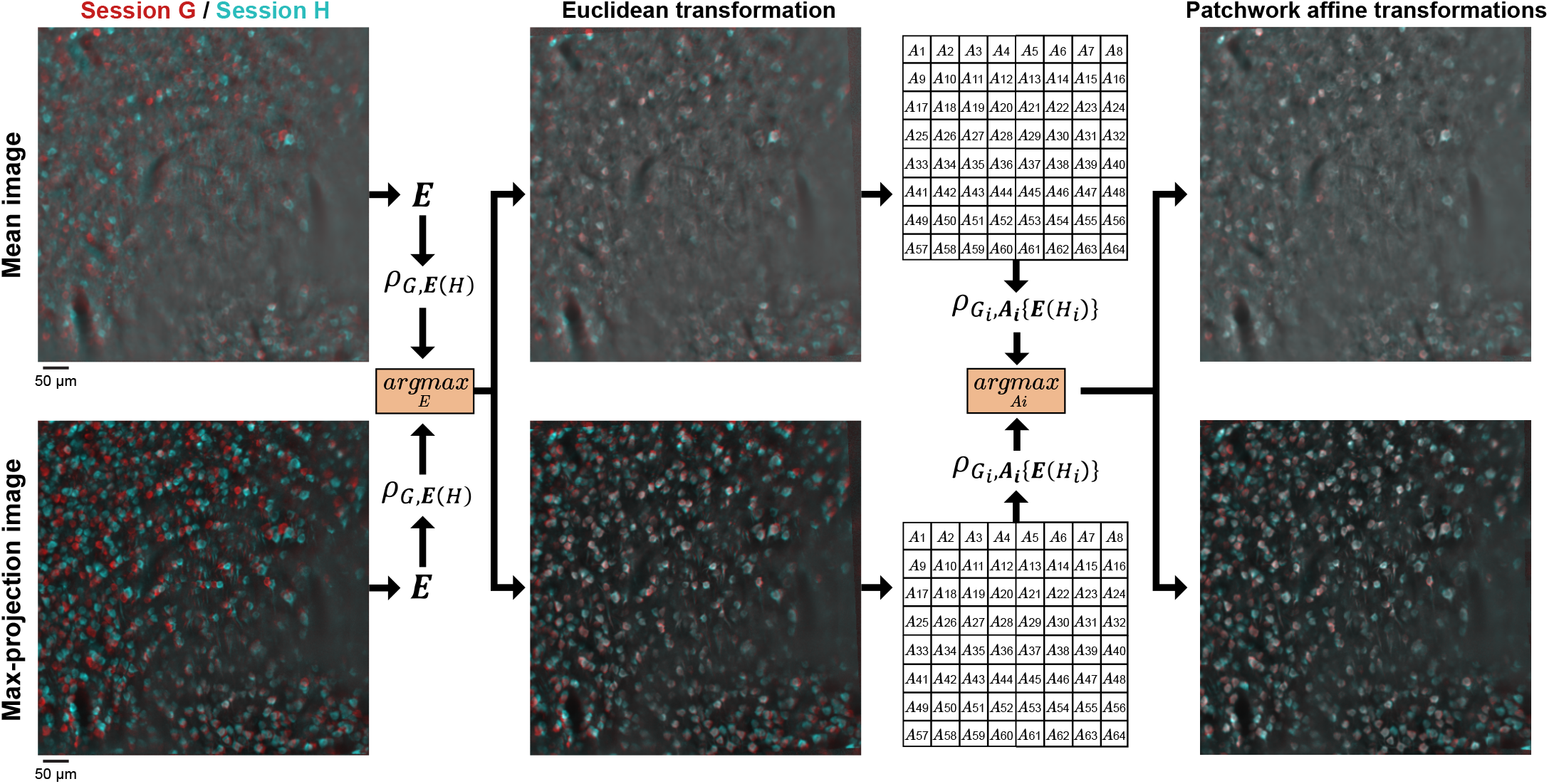
PatchWarp algorithm for fully automac across-session registration. Registration between different imaging sessions for longitudinal analyses (Red: Session G images, Cyan: Session H images). Mean and max-projection images were used as the inputs to the algorithm. The first step finds the optimal Euclidean transformation that corrects large displacements with rigidity (translation and rotation).The second step finds the optimal affine transformations for small patches to correct non-uniform distortions between the 2 image sessions. The transformations are separately estimated for mean and max-projection images, but only one transformation with larger ECC between G and transformed 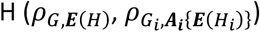 is selected at each step. The obtained transformations can be used to transform any images from session H to the session G coordinate for longitudinal neural activity analyses.

The first step finds the optimal Euclidean transformation that maximizes ECC between the session G summary images and the transformed session H summary images. The whole FOVs are used without splitting them into patches in this first step. The Euclidean transformation is a special case of affine transformation with reduced number of free parameters (6 and 3 parameters in affine and Euclidean transformation matrices, respectively), and the transformation is limited to translation and rotation. Unlike the rigid motion correction step of the within-session PatchWarp processing where the corrections were limited to translations, we use Euclidean transformations here because mismatch in the rotation angles of the imaging FOVs is common across different imaging sessions. To ensure the robust estimation of *E* even in the presence of a large displacement between the 2 imaging sessions, we took the pyramid-based approach by sequentially updating the transformation matrix from the upper level pyramids at the gradient-based parameter estimation step (see Methods). The transformation that maximizes ECC between the session G images and the transformed session H images was obtained. We locally normalized the intensity of the input summary images before calculating ECC to focus on neuronal structures instead of calcium levels, similarly to the within-session warp correction method (Figure S1). We use 2 types of summary images (mean and max-projection images) and obtain 2 transformation matrices. We select the one of them that results in a higher ECC, and apply it to both of the session H summary images.

The second step finds the optimal affine transformations that maximize ECCs between the session G patches and the transformed session H patches. Large displacements are already corrected by the first Euclidean transformation, so the goal of this second set of transformations is to correct the non-uniform image distortions between the 2 imaging sessions. The affine transformation matrices with 6 parameters can transform image patches by shearing and scaling in addition to translation and rotation. By selecting the transformation matrix with larger ECC for each patch from the mean and the max-projection image pairs, we can optimally correct non-uniform image distortions even when a patch lacks sufficient landmarks in one of the summary images.

The two-step PatchWarp algorithm for across-session registration provides us with the transformation matrices that can optimally transform the session H images to the session G coordinate. The obtained transformation can be applied to any images in the session H coordinate (e.g. individual frames, summary images such as mean and max-projection, correlation map, ROI mask), and the transformed images can be directly compared with the session G images for longitudinal activity analyses. We provide functions to calculate the transformation and apply the transformation to other images in our software.

## Discussion

Removal of activity-independent artifacts from calcium imaging data is critical to accurately interpret data. For example, rigid motion artifacts may be misinterpreted as movement-related neural activity, and slow image distortions may be misinterpreted as slow activity changes over time. These artifacts can have profound impacts on the neural activity analyses and potentially lead to incorrect scientific conclusions. Here, we developed a novel algorithm, PatchWarp, that robustly corrects both the rigid motion artifacts and slow image distortions. We publicly share the software at https://github.com/ryhattori/PatchWarp for the community.

The warp correction step of PatchWarp splits a FOV into smaller patches, and optimal affine transformation is estimated for each patch. Two existing and popular motion correction algorithms for 2-photon calcium imaging data, Suite2p and CaImAn (NoRMCorre), offer non-rigid registration algorithms that similarly split a FOV into small patches. However, the transformation of each patch in these published methods is limited to translational shifts. Although translational shifts of many small patches can approximate shearing transformation, the accuracy is suboptimal. The rigidity on each patch limits the capacity of the fine-scale distortion corrections unless each patch is very small, and the patch cannot be too small given the limited number of spatial landmarks. In contrast, PatchWarp applies an affine transformation to each patch. Affine transformation can transform each patch not only by translation but also rotation, scaling, and shearing. Utilizing the diverse transformational capacity of the affine transformation, we achieved a higher registration accuracy than Suite2p and CaImAn (NoRMCorre) (Figure 5).

Although PatchWarp is robust in correcting rigid motion artifacts and slow image distortions, we still observed residual motion artifacts in some cases especially when the neuronal structures are very close to the midline sinus. The residual motion artifacts are due to fast dilation of the midline sinus which occasionally happens in behaving mice. The dilation of sinus introduces fast image distortions. Since PatchWarp estimates the affine transformations using mean of multiple frames for the robustness and the computational efficiency, it mostly focuses on correcting slow image distortions. Although users can apply PatchWarp to correct fast image distortions by using smaller number of frames to make the mean image to register to the template image, the accuracy of warp correction could be compromised if each mean image does not contain sufficient neuronal structures with visible calcium signals. The dynamic nature of calcium signals introduces uncertainty about the availability of sufficient signals on each mean image. If users need to correct fast image distortions, one potential solution is to co-express stable red fluorescence indicators (e.g. tdTomato (Drobizhev et al., 2011; Shaner et al., 2004)) and use the signals for the registration instead of using the dynamic calcium signals. Alternatively, experimenters should always try to minimize the contributions of the sinus dilations by applying enough pressure when they implant a glass window onto the brain or avoid selecting a FOV near the midline sinus.

We also developed a PatchWarp approach for fully automated across-session image registrations. Registration across images from different imaging sessions is necessary to longitudinally analyze how individual neurons or neuronal structures change activity patterns over days. Since our method uses only the summary images from each session, the estimation of the optimal transformation matrices can be obtained instantly with only 3 to 6 seconds on standard desktop computers. By applying the obtained transformations, we can transform images to the coordinate of the other session for longitudinal neural activity analyses.

## Supplementary figure and video legends

**Supplementary figure 1. Normalization of mean images by their local intensity**.

**A)** Steps for making a blurred image (local intensity map) which will be used to normalize a mean image. First, the original mean image was convolved by a circular filter with a radius of 32 pixels. The edge pixels are dim due to zero-paddings. To correct the zero-padding effects, we apply the same convolution to all-ones matrix with the same pixel number as the mean image. The resulting convolved all-ones matrix image also exhibits dim intensity near the edges due to zero-paddings. Therefore, division of the convolved mean image by the convolved all-ones matrix results in a blurred image without zero-padding artifacts. **B)** Steps to make an image with local intensity normalization. The intensity of each pixel in the blurred image from **(A)** reflects the local intensity near the pixel. Therefore, simple division of the original mean image by the blurred image normalizes each pixel intensity by its surrounding intensity. To return the intensity value back to the original scale, the division is multiplied by the mean intensity of the original mean image.

**Supplementary figure 2. mCM with consistent template images for comparisons between pre- and post-rigid corrections**.

**A-C)** Mean correlation between the mean of either raw frames (Raw template) or rigidly corrected frames (Rigid template) and individual frames. mCM consistently increases after rigid motion corrections, regardless of which template image is used to calculate the correlation. Therefore, the difference in self-mCM in Figure 3B is not due to the different template images for Raw and Rigid conditions.

**Supplementary figure 3. mCM with consistent template images for comparisons between pre- and post-warp corrections**.

**A-C)** Mean correlation between the mean of either only rigidly corrected frames (Rigid template) or distortion-corrected frames (Warp template) and individual frames. mCM consistently increases after warp corrections, regardless of which template image is used to calculate the correlation. Therefore, the difference in self-mCM in Figure 4B is not due to the different template images for Rigid and Warp conditions.

**Supplementary video 1. Example image frames before and after the rigid motion corrections**. Image frames before (*Left*) and after (*Right*) rigid motion corrections. Frames that were acquired at the frame rate of 29 Hz from an example ALM imaging session are shown for 17.2 sec. Note that the video is sped up. The time stamp is shown on the bottom left.

**Supplementary video 2. Example images with only rigid-motion corrections or with both rigid and warp corrections**.

Images with only rigid-motion corrections (*Left*) or with both rigid and warp corrections (*Right*) from an example pM2 imaging session that lasted for 118 minutes. Images were temporally smoothed by a Gaussian filter (σ = 11,600 frames). Slow distortions are corrected by the warp corrections. Note that the video is sped up. The time stamp is shown on the bottom left.

## Methods

### Animals

Mice were originally obtained from the Jackson Laboratory (CaMKIIa-tTA: B6;CBA-Tg(Camk2a-tTA)1Mmay/J [JAX 003010]; tetO-GCaMP6s: B6;DBA-Tg(tetO-GCaMP6s)2Niell/J [JAX 024742]; B6;129S6-Chattm2(cre)Lowl/J [JAX:006410]). All animals were housed in disposable plastic cages with standard bedding in a room on a reversed light cycle (12 h/12 h). All procedures were performed following protocols approved by the UCSD Institutional Animal Care and Use Committee and guidelines of the National Institute of Health.

### Surgery

Surgical procedures were performed as previously described (Hattori et al., 2019). Mice were continuously anesthetized with 1-2% isoflurane during surgery. We subcutaneously injected dexamethasone (2mg/kg). We exposed the dorsal skull, removed the connective tissue on the skull surface using a razor blade, and performed craniotomy. For axon imaging experiments, ∼1 μL of AAV-hSyn-FLEX-axon-GCaMP6s virus (Broussard et al., 2018) was unilaterally injected in the basal forebrain of ChAT-Cre mice (∼0.3 mm posterior and ∼1.7 mm lateral from bregma, 5.0 mm deep from the brain surface). After the injection, a glass window was secured on the edges of the remaining dorsal skull using 3M Vetbond (WPI), followed by cyanoacrylate glue and dental acrylic cement (Lang Dental). For GRIN lens imaging experiments, we first aspirated the cortex above the target coordinate up to 1.0 mm depth using a blunt end 30G needle (0.312 mm O.D., SAI Infusion Technologies). Then, a GRIN lens (500 µm diameter; Inscopix, GLP-0561) was unilaterally implanted above the deep layer of lateral OFC ((∼2.45 mm lateral and ∼2.6 mm anterior from bregma, 1.5 mm deep from the brain surface). The implanted GRIN lens was secured using 3 M Vetbond (WPI) on the skull, followed by cyanoacrylate glue and dental acrylic cement (Lang Dental). After the implantation of either a glass window or a GRIN lens, a custom-built metal head-bar was secured on the skull above the cerebellum using cyanoacrylate glue and dental cement. Mice were subcutaneously injected with Buprenorphine (0.1 mg/kg) and Baytril (10 mg/kg) after surgery.

### Two-photon calcium imaging

Neural calcium signals were recorded using two-photon microscopes (B-SCOPE, Thorlabs) with a 16×, 0.8 NA water immersion objective lens (Nikon) and 925 nm lasers (Ti-Sapphire laser, Newport) from head-fixed behaving mice. ScanImage (Vidrio Technologies) running on MATLAB (MathWorks) was used for image acquisitions. Images (512 × 512 pixels) were continuously recorded at ∼29 Hz during each imaging session. Cell body imaging sessions were recorded for 1.5-3.5 hours while axon imaging sessions were recorded for ∼20 minutes. Imaging was performed at superficial layers of cortex for glass window imaging, and GRIN lens imaging was performed at deep layers of OFC. The central coordinates of FOVs from bregma were 1.7 mm lateral and 2.25 mm anterior for ALM, 0.4 mm lateral and 0.5 mm anterior for pM2, 0.4 mm lateral and 2 mm posterior for RSC, 1.7 mm lateral and 2 mm posterior for PPC, 1.8 mm lateral and 0.75 mm posterior for S1, 2.5 mm lateral and 3.25 mm posterior for V1.

### Template re-estimation for rigid motion correction

Rigid motion artifacts during calcium imaging were corrected by registering individual frames to a template image. The template image was iteratively updated during the motion correction process. This iterative re-estimations of the template image improve the robustness of motion correction performance, especially in the presence of slow image distortion over time. First, we created the 1^st^ template image (*T*_1_) by averaging the middle 2,500 frames of an imaging session. Then, we registered the 2,500 frames to the template *T*_1_ and updated the *T*_1_ by averaging the motion-corrected 2,500 frames 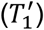. Each imaging session was temporally split into 5 blocks, and 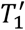 was used as the template image to register frames in the 3^rd^ block. After registering all frames in the 3^rd^ block, we created 2 new template images by averaging the first and the last 2,500 motion-corrected frames in the 3^rd^ block (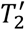 and 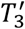, respectively). The template 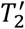 was used to register all frames in the 2^nd^ block, and the template 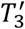 was used to register all frames in the 4^th^ block. Again, we created new template images by averaging the first 2,500 frames of the 2^nd^ block 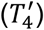 and averaging the last 2,500 frames of the 4^th^ block 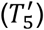. The template image 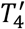 was used to register frames in the 1^st^ block, and the template image 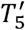 was used to register frames in the 5^th^ block.

### Registration algorithm for rigid motion correction

Individual frames were shifted along x and y directions such that the correlation between each frame and a template image is maximized. To estimate the best translational shift of each frame, pyramid method (Adelson et al., 1984) and hill climbing algorithm were combined as reported previously (Mitani and Komiyama, 2018). First, both the template image and individual frames were downscaled to the level 3 pyramid representation (1/8 downscaling factor). Hill climbing algorithm finds the best translation that maximizes the correlation between each frame and the template. Then, the algorithm moves to the level 2 pyramid representation (1/4 downscaling factor) and uses the translation from the level 3 as the initial estimate of the translational shift to obtain the best shift at the level 2. It repeats the same procedure at the level 1 pyramid representation (1/2 downscaling factor) and lastly at the level 0 (original image resolution). The registration was performed in subpixel accuracy by fitting a quadratic function and interpolation. The pixel intensities of both the template image and individual frames were rank transformed before running the pyramid-based hill climbing algorithm.

### Registration algorithm for slow distortion correction

We performed affine transformations of images to correct slow image distortion. However, image distortions in calcium imaging data are usually not uniform across the FOV, making it impossible to correct the distortion by a single affine transformation matrix. To correct non-uniform distortions, we developed a novel patchwork approach of affine transformation. Our approach assigns different affine transformation matrices to different subfields of an image, and the transformed subfields are stitched together. This patchwork approach efficiently and robustly corrects non-uniform image distortions in calcium imaging data.

We split each FOV of the size H × W (pixels) into M × M square subfields (patches). We used M = 8 for cellular resolution imaging and used M = 12 or 15 for axon imaging. We used larger M for axon imaging because axons were more dynamic and the movements were independent from each other. Each patch had 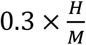 overlapping pixels with its vertically adjacent patches and 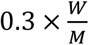 overlapping pixels with its horizontally adjacent patches. After affine transformation of each patch, all patches were stitched together by averaging the intensity of the overlapping pixels.

The template image for each patch was created by averaging the middle 5,500 frames of an imaging session. We then temporally downsampled all frames by non-overlapping moving averaging of 500 frames. Optimal affine transformation for each patch was obtained for each of the downsampled frames. This downsampling significantly decreases the computation time and did not affect our results because the slow image distortions were negligible within 500 frames (∼17 sec, ∼29 Hz).

An affine transformation matrix for a 2D image is given by the following 3 × 3 matrix with 6 parameters;

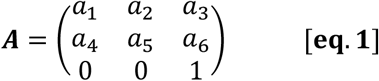

The relationships between the original coordinate (*x, y*) and the transformed coordinate (*x*′, *y*′) is given by

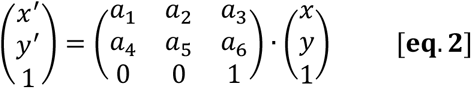

To find the optimal affine transformation for correcting image distortion of each patch, we used a gradient-based algorithm as described previously (Evangelidis and Psarakis, 2008). Our objective is to minimize the difference between a template image patch and a warped image patch, but we want to focus on the geometric difference while ignoring brightness difference because brightness changes across frames on calcium imaging data. We can define the loss function of the gradient-based algorithm as follows;

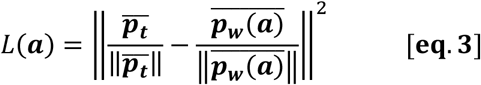

where ‖ · ‖ denotes L2 norm, ***a*** = [*a*_1_, *a*_2_, *a*_3_, *a*_4_, *a*_5_, *a*_6_] is a vector of parameters for the affine transformation, ***p***_***t***_ is a vector with intensity of all pixels in the template patch, ***p***_***w***_(***a***) is a vector with intensity of all pixels in the patch that was warped by an affine transformation matrix with the parameters 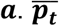 and 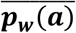 denote the zero-centered vectors of ***p***_***t***_ and ***p***_***w***_(***a***), respectively. The pixel intensity normalizations of both the template and warped images in [eq. 3] ensure that the loss function is invariant to changes in bias, gain, brightness and contrast. Thus, we can selectively minimize geometric difference using the loss function. We can equivalently express the [eq. 3] as follows;

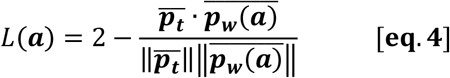

Therefore, minimizing *L*(***a***) is equivalent to maximizing the following index;

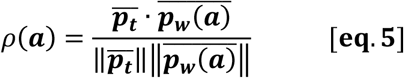

where 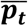 and 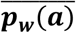 are zero-centered vectors of ***p***_***t***_ and ***p***_***w***_ (***a***). This index is called as enhanced correlation coefficient (ECC). We estimated the optimal affine transformation matrix that maximizes ECC using a gradient-based approach (Evangelidis and Psarakis, 2008; Psarakis and Evangelidis, 2005).

To further improve the robustness and accuracy of the gradient-based algorithm, we processed both input images to normalize the temporal dynamics of calcium signals. First, we obtained blurred images by performing 2D convolution using a circular kernel (32 pixel radius) as follows;

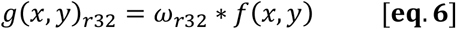

where * denotes convolution operation, *g*(*x, y*) is the original image, *ω*_*r*32_ is a circular kernel with 32 pixel radius, and *g*(*x, y*)_*r*32_ is the blurred image. The pixel intensity near the edges of the blurred images are low due to zero-padding. To compensate for the zero-padding effect, we normalized the blurred images as follows;

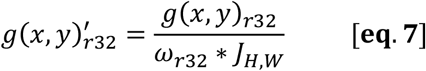

where *J*_*H,W*_ is an all-ones matrix where every element is equal to 1. Using the blurred images, we normalized the intensity of the original image as follows;

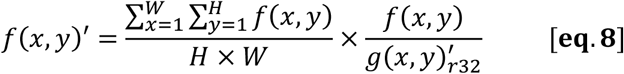

This intensity normalization enhances the contrast between dim neuronal structures and their local surrounding areas (radius of 32 pixels) on each frame, which helps the gradient-based algorithm because the changes in neuronal calcium-dependent fluorescence intensity across frames are normalized by this intensity normalization.

After deriving the optimal affine transformation matrix for each image patch, we transformed each patch by subpixel bilinear interpolation such that the x-y coordinate is transformed following the optimal affine transformation. The transformed patches were stitched together by averaging the intensity of overlapping pixels from adjacent patches to obtain the transformed full-FOV image.

### Registration between different imaging sessions

We developed a robust automated approach to register images between different imaging sessions for longitudinal neural activity analyses. The inputs to the algorithm are summary images from each imaging session. It accepts any summary images such as the mean image (intensity is averaged across frames for each pixel), the max-projection image (max-projection of intensity from all frames for each pixel), the standard deviation image (standard deviation of intensity across frames for each pixel), and the correlation image (average correlation across frames between each pixel and its neighbors). Here, we use mean and max-projection images as the example inputs. We denote the mean and max-projection images of session G and session H as *f*(*x, y*)_*mean,G*_, *f*(*x, y*)_*mean,H*_, *f*(*x, y*)_*max,G*_, and *f*(*x, y*)_*max,H*_, respectively. First, we locally normalize the intensity of each summary image using [eq. 8] and obtain 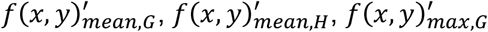, and 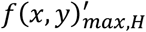. Our goal is to obtain the optimal transformation matrices for the session H summary images such that transformed summary images of session H match the summary images of session G. We obtain the optimal transformation for the session H with a 2-step process.

The first step finds the optimal Euclidean transformation that maximizes ECC between the session G summary images and the transformed session H summary images. The whole FOVs are used without splitting them into patches in this first step. The Euclidean transformation matrix for a 2D image is given by the following 3 × 3 matrix with 3 parameters;

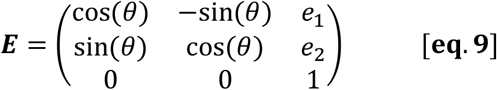

The Euclidean transformation matrix is a special case of an affine transformation matrix, and the transformation is limited to translation (*e*_1_ for x-shift, *e*_2_ for y-shift) and rotation (*θ*). The reduced number of parameters (6 parameters in ***A***, 3 parameters in ***E***) makes the estimation of the best translational and rotational shifts more robust than when the affine transformation matrix with full 6 parameters are used.

To ensure the robust estimation of ***E*** even in the presence of a large displacement between the 2 imaging sessions, we took the pyramid-based approach. First, intensity-normalized images 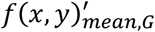 and 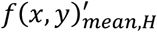 were downscaled to the level 3 pyramid representation (1/8 downscaling factor). A gradient-based algorithm finds the best ***E*** that maximizes ECC [eq. 5] between the session G image 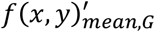 and the transformed session H image 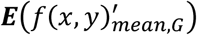 using the level 3 pyramid representations. Then, the algorithm moves to the level 2 pyramid representation (1/4 downscaling factor) and uses ***E*** from the level 3 as the initial estimate to obtain the best ***E*** at the level 2. It repeats the same procedure at the level 1 pyramid representation (1/2 downscaling factor) and lastly at the level 0 (original image resolution). We performed the pyramid-based approach for both the mean and max-projection image pairs to obtain the optimal Euclidean transformation matrices ***E***_***mean***_ and ***E***_***max***_. We then selects the Euclidean transformation matrix that gives the largest ECC as follows;

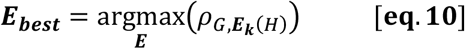

where ***E***_***k***_ is either ***E***_***mean***_ or ***E***_***mean***_, and 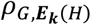 denotes ECC between the summary image from the session G and the transformed summary image from the session H. We used ***E***_***best***_ to transform 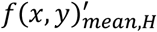 and 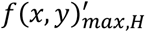 to 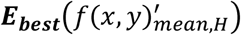 and 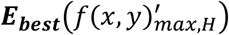, respectively, by subpixel bilinear interpolation.

The above Euclidean transformation registers the 2 imaging sessions using translational and rotational shifts. However, these rigid shifts are not sufficient for optimal registration due to slight differences in focal planes (e.g. rotation angle of the objective lens, z-depth of the focal plane) and deformation of the imaged brain tissue between the 2 imaging sessions. Therefore, the next second step aims at solving this non-rigid registration problem using the method we described in the previous section (“*Registration algorithm for slow distortion correction”*). We split 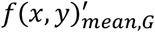 and 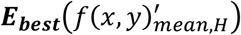 into M × M square patches and find the optimal affine transformation for each patch. We use affine transformation matrices with full 6 parameters [eq. 1] to allow not only translation and rotation but also scaling and shearing for the registration. We do not need to implement pyramid method in this second step because the first Euclidean transformation already corrected large displacements.

We estimated the optimal affine transformations for all patches of both the mean and max-projection image pairs. We denote the optimal affine transformation matrices for *i*^*th*^ patch by ***A***_***i***,***mean***_ and ***A***_***i***,***max***_. We then selects the affine transformation matrix that gives the largest ECC for each patch as follows;

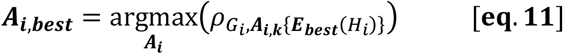

where *G*_*i*_ is the *i*^*th*^ patch from session G, *H*_*i*_ is the *i*^*th*^ patch from session H, ***A***_***i***,***k***_ is either ***A***_***i***,***mean***_ or ***A***_***i***,***max***_, and 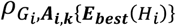 denotes ECC between the *i*^*th*^ patch from the session G and the transformed *i*^*th*^ patch from the session H. We can use ***E***_***best***_ and ***A***_***i***,***best***_ to transform *i*^*th*^ patches 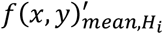 and 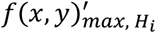 to 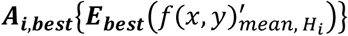 and 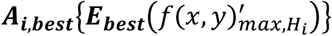, respectively, by subpixel bilinear interpolation. We repeat these steps for all patches, so the number of affine transformation matrices that we obtain is M × M. In total, we obtain (M × M + 1) transformation matrices (including both ***E***_***best***_ and ***A***_***i***,***best***_) for a pair of imaging sessions which we want to register. Using these transformation matrices, we can transform individual imaging frames or other summary images (e.g. mean image, max-projection image, correlation image, standard deviation image, ROI mask image) from session H to session G coordinate for longitudinal neural activity analyses.

### Data analysis software and library

PatchWarp software was written in MATLAB. We used MATLAB and Python3.7 for data analyses. The original codes for pyramid-based hill climbing algorithm (Mitani and Komiyama, 2018) and gradient-based ECC maximization algorithm (Evangelidis and Psarakis, 2008) are available at https://github.com/amitani/matlab_motion_correct and https://www.mathworks.com/matlabcentral/fileexchange/27253, respectively.

## Data and code availability

All datasets and analysis codes used in this study are available from the corresponding author upon reasonable request.

## Code availability

PatchWarp software is available at https://github.com/ryhattori/PatchWarp.

## Acknowledgements

We thank Akinori Mitani for providing codes for pyramid-based hill climbing algorithm and consultation at early stage of this project, Chi Ren for providing axon imaging data, and the members of the Komiyama lab for discussions and feedbacks. R.H. was supported by the Uehara Memorial Foundation Postdoctoral Fellowship, JSPS Postdoctoral Fellowship for Research Abroad, the Research Grant from the Kanae Foundation for the Promotion of Medical Science, and The Warren Alpert Distinguished Scholar Awards. T.K. was supported by grants from NIH (R01 NS091010, R01 EY025349, R01 DC014690, and P30 EY022589), NSF (1940181), and David & Lucile Packard Foundation.

## Author contributions

RH conceived the methods, developed the software, performed analyses, and collected cell body imaging data. RH and TK wrote the paper.

## Notes

### Competing Interest Statement

The authors have declared no competing interest.

