## Supplementary Figures for "PatchWarp: Corrections of non-uniform image distortions in two-photon calcium imaging data by patchwork affine transformations"

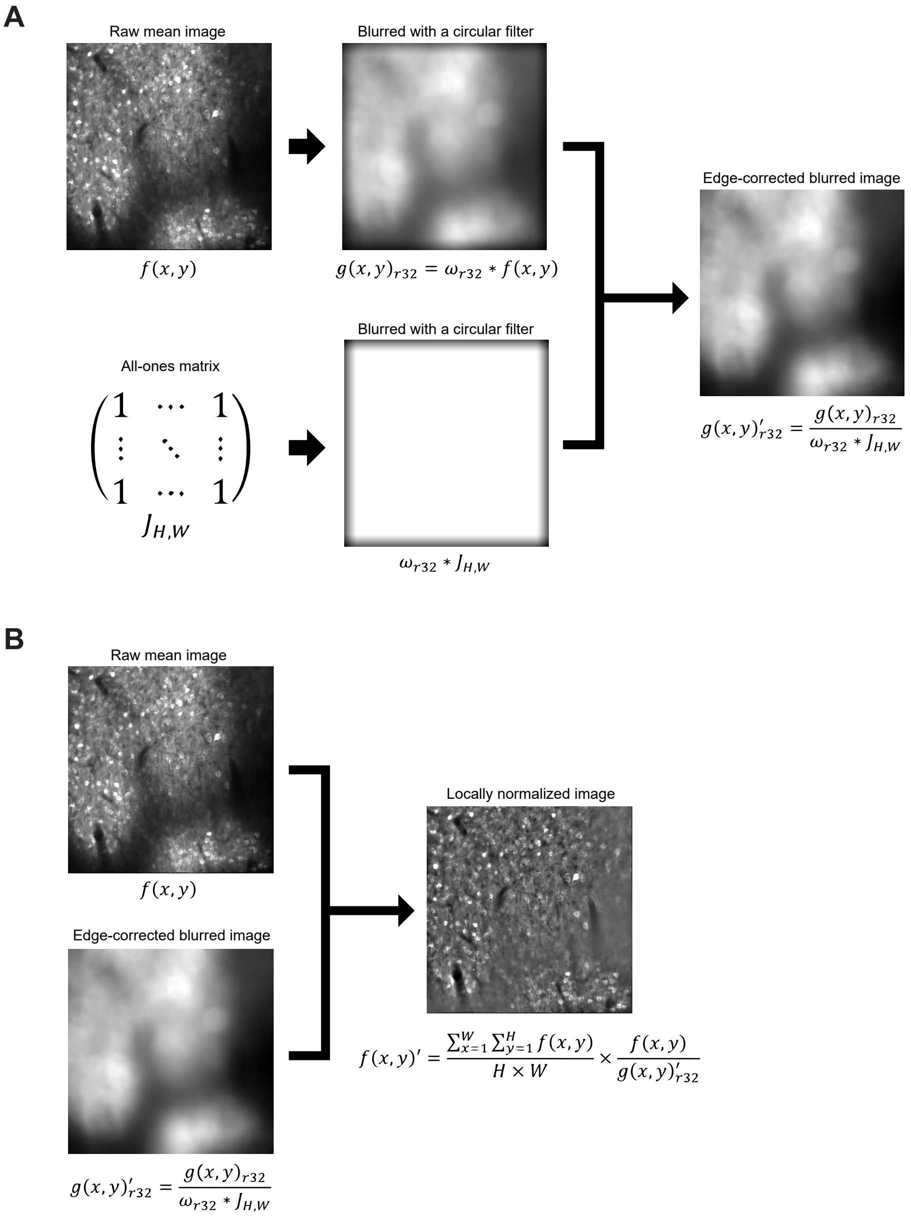


**Supplementary figure 1. Normalization of mean images by their local intensity.**
**A)** Steps for making a blurred image (local intensity map) which will be used to normalize a mean image. First, the original mean image was convolved by a circular filter with a radius of 32 pixels. The edge pixels are dim due to zero-paddings. To correct the zero-padding effects, we apply the same convolution to all-ones matrix with the same pixel number as the mean image. The resulting convolved all-ones matrix image also exhibits dim intensity near the edges due to zero-paddings. Therefore, division of the convolved mean image by the convolved all-ones matrix results in a blurred image without zero-padding artifacts. **B)** Steps to make an image with local intensity normalization. The intensity of each pixel in the blurred image from **(A)** reflects the local intensity near the pixel. Therefore, simple division of the original mean image by the blurred image normalizes each pixel intensity by its surrounding intensity. To return the intensity value back to the original scale, the division is multiplied by the mean intensity of the original mean image.

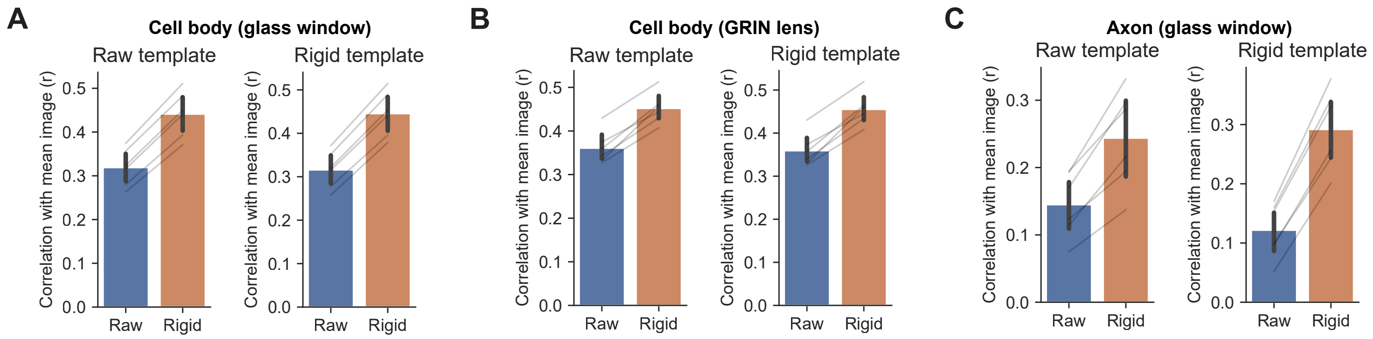


**Supplementary figure 2. mCM with consistent template images for comparisons between pre- and post-rigid corrections.**

**A-C)** Mean correlation between the mean of either raw frames (Raw template) or rigidly corrected frames (Rigid template) and individual frames. mCM consistently increases after rigid motion corrections, regardless of which template image is used to calculate the correlation. Therefore, the difference in self-mCM in Figure 3B is not due to the different template images for Raw and Rigid conditions.

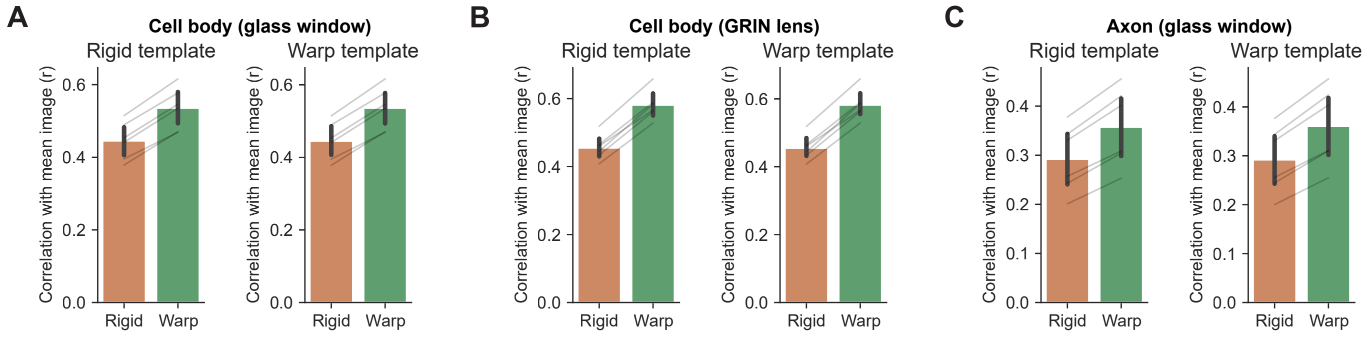


**Supplementary figure 3. mCM with consistent template images for comparisons between pre- and post-warp corrections.**

**A-C)** Mean correlation between the mean of either only rigidly corrected frames (Rigid template) or distortion-corrected frames (Warp template) and individual frames. mCM consistently increases after warp corrections, regardless of which template image is used to calculate the correlation. Therefore, the difference in self-mCM in Figure 4B is not due to the different template images for Rigid and Warp conditions.
